# Thermal Stimulus Task fMRI in the Cervical Spinal Cord at 7 Tesla

**DOI:** 10.1101/2023.01.31.526451

**Authors:** Alan C. Seifert, Junqian Xu, Yazhuo Kong, Falk Eippert, Karla L. Miller, Irene Tracey, S. Johanna Vannesjo

**Author notes:** Corresponding author: Alan C. Seifert: Address: One Gustave L. Levy Place, Box 1234, New York, NY 10029. **Submitted to: Human Brain Mapping**. Preliminary results from this manuscript have previously been presented in the following conference abstracts: Seifert AC, Vannesjo SJ. Spatial Specificity of BOLD Signal in the Spinal Cord at 7T Using a Noxious Thermal Stimulus. Proceedings of the International Society for Magnetic Resonance in Medicine 2020:1168; Seifert AC, Vannesjo SJ. Comparison of Physiological Noise Models for Thermal Stimulus fMRI in the Cervical Spinal Cord at 7T. Proceedings of the International Society for Magnetic Resonance in Medicine 2022:1462.

## Abstract

**Purpose:** Although functional MRI is widely applied in the brain, fMRI of the spinal cord is more technically demanding. Proximity to the vertebral column and lungs results in strong spatial inhomogeneity and temporal fluctuations in B_0_. Increasing field strength enables higher spatial resolution and improved sensitivity to BOLD signal, but amplifies the effects of B_0_ inhomogeneity. In this work, we present the first task fMRI in the spinal cord at 7 T. Further, we compare the performance of single-shot and multi-shot 2D EPI protocols, which differ in sensitivity to spatial and temporal B_0_ inhomogeneity.

**Methods:** The cervical spinal cords of 11 healthy volunteers were scanned at 7 T using single-shot 2D EPI at 0.75 mm in-plane resolution and multi-shot 2D EPI at 0.75 and 0.6 mm in-plane resolutions. All protocols used 3 mm slice thickness. For each protocol, the BOLD response to thirteen 10-second noxious thermal stimuli applied to the right thumb was acquired in a 10-minute fMRI run. Image quality, temporal SNR, and BOLD activation (percent signal change and z-stat) at both individual-and group-level were evaluated between the protocols.

**Results:** Temporal SNR was highest in single-shot and multi-shot 0.75 mm protocols. In group-level analyses, activation clusters appeared in all protocols in the ipsilateral dorsal quadrant at the expected C6 neurological level. In individual-level analyses, activation clusters at the expected level were detected in some, but not all subjects and protocols. Single-shot 0.75 mm generally produced the highest mean z-statistic, while multi-shot 0.60 mm produced the best-localized activation clusters and the least geometric distortion. Larger than expected within-subject segmental variation of BOLD activation along the cord was observed.

**Conclusion:** Group-level sensory task fMRI of the cervical spinal cord is feasible at 7 T with single-shot or multi-shot EPI. The best choice of protocol will likely depend on the relative importance of sensitivity to activation versus spatial localization of activation for a given experiment.

**Highlights:**

First stimulus task fMRI results in the spinal cord at 7 T
Single-shot 0.75 mm 2D EPI produced the highest mean z-statistic
Multi-shot 0.60 mm 2D EPI provided the best-localized activation and least distortion

## Introduction

The spinal cord houses many scientifically and clinically important neural circuits, such as central pattern generators related to locomotion^1^, sensorimotor reflex arcs, and circuits involved in the modulation of nociception^2^. Although functional MRI is a well-established and widely-applied imaging modality to probe neural activity in the brain, fMRI of the spinal cord is more technically demanding.

The adult human spinal cord is approximately 15 mm in diameter at its widest point, with gray matter structures only approximately 1-4 mm wide. High spatial resolution is therefore required to image the blood oxygenation level-dependent (BOLD) signal in the spinal cord. In addition to its small size, several additional factors complicate fMRI of the spinal cord. The spinal cord is directly adjacent to the vertebral column, which is composed of alternating vertebral bones and soft tissues that differ in magnetic susceptibility, introducing distortions in the B_0_ field. These spatially periodic B_0_ inhomogeneities in the spinal cord cannot be adequately cancelled by a counter-field generated by the scanner’s shim coils, leaving residual focal B_0_ inhomogeneities near the intervertebral junctions. The spinal cord is also near the lungs, whose volume and concentration of paramagnetic O_2_ varies over time due to respiration, also introducing B_0_ distortions. The resulting temporal fluctuations in B_0_ in the spinal cord^3^ produce frequency variation that can exceed 100 Hz at C7^4^. These spatially and temporally varying B_0_ inhomogeneities produce geometric distortion and signal loss, as well as apparent motion between volumes in the time series. Finally, pulsatile rostral-caudal flow of cerebrospinal fluid (CSF) over the cardiac cycle, and intermittent swallowing and motion, can also contribute nuisance variations to fMRI data that are more extreme than in the cerebrum^5,6^.

Increasing the static magnetic field (B_0_) improves signal to noise ratio (SNR), which translates to higher achievable spatial resolution. Increased B_0_ also increases the BOLD signal acquired by gradient-echo (GRE) EPI, leading to better sensitivity and spatial specificity^7,8^. These advantages motivate the use of 7 T fMRI to investigate circuits in the spinal cord. However, several difficulties are magnified and become significant at 7 T or higher field strengths. As field strength increases, susceptibility-mediated effects are magnified. Hence, although SNR and BOLD signal are enhanced, the geometric distortions and signal loss associated with B_0_ inhomogeneity arising from the lungs and vertebral column are also intensified^9,10^. Most 7 T scanners also lack a body RF transmit coil, and standard head RF coils do not cover the cervical spinal cord, which necessitates specialized RF hardware for the spinal cord that remains only accessible at a handful of research centers^11^.

These challenges have hampered the development of spinal cord fMRI at 7 T, despite the potential advantages of higher field strength. To date there are only three studies of spinal cord fMRI at 7 T^12–14^ all of which are resting-state fMRI studies investigating connectivity between gray matter regions in the absence of external stimuli or motor tasks. A significant motivation for the development of spinal cord fMRI methods, however, is to study pain processing in healthy and pathological states^15–20^. This involves measuring the BOLD response to external painful stimuli. There is thus a need for investigating the potential of 7 T for task-based spinal cord fMRI.

One important step towards fully realizing the potential benefits of 7 T is to establish suitable acquisition sequences and protocols that yield high BOLD sensitivity while being sufficiently robust against the challenges posed by static and dynamic B_0_ inhomogeneities. At lower field strengths, various sequences have been investigated for spinal cord fMRI. Single-shot echo-planar imaging (EPI) is currently the dominant method for fMRI in the human spinal cord, as in the brain. The initial motor task fMRI studies in the cervical spinal cord, however, used a variety of pulse sequences, including fast low-angle shot (FLASH) sequences at 1.5 T^21^ and 3 T^22^, multi-shot gradient echo (GRE) EPI at 1.5 T^23^, spin-echo EPI at 1.5 T^24^, and single-shot fast spin echo at 1.5 T^25,26^. Spin-echo-based pulse sequences continued to be used for fMRI in the spinal cord at 1.5 T based on signal enhancement by extravascular water protons (SEEP) contrast for many years^27,28^. Work on BOLD fMRI in the spinal cord by multiple groups over the next several years focused mainly on 2D single-shot^29–35^ and 3D multi-shot^36^ GRE EPI at 1.5 T and 3 T.

Most relevant to the present work, others have previously implemented 2D single-shot EPI in combination with noxious thermal stimulation to study cognitive manipulation of pain^17,37^ the effects of noxious thermal stimulation on functional connectivity in the cervical spinal cord^38^, and several have developed and implemented methods for simultaneous brain and spinal cord MRI^39–41^ and utilized them in studies of, for example, the fundamental organization of resting-state networks spanning the brain and the cervical spinal cord^42,43^ and the interaction between brain and spinal cord regions on sensitivity to pain^19,44^.

Single-shot EPI sequences are heavily affected by geometric distortion and signal loss caused by static B_0_ inhomogeneity. Multi-shot EPI sequences have strong potential to reduce these artifacts by shortening the echo train length and echo time. In the previous 7 T resting-state studies, highly segmented multi-shot 3D EPI with very short echo train lengths was used^12,13,45^, yielding images largely uncorrupted by signal loss and distortion near intervertebral junctions. However, the transition from single-shot to multi-shot EPI entails a penalty in volume acquisition time (VAT), which can reduce statistical power to detect to BOLD activation by reducing the number of observation time points for a given scan duration. Multi-shot EPI is also more vulnerable to dynamic B_0_ field fluctuations, resulting in variable ghosting between volumes in the time series, which can reduce the temporal SNR^46^. In single-shot EPI, on the other hand, time-variable B_0_ fields (at the respiratory time-scale) largely translate into apparent motion in the images, which can be addressed by retrospective motion correction. Considering the relative advantages and disadvantages of single-shot vs. multi-shot EPI sequences, it is not apparent which is more suitable for fMRI of the spinal cord at ultra-high field.

In this work, we present the first stimulus task fMRI in the cervical spinal cord at 7 T and compare the performance of one single-shot 2D EPI and two multi-shot 2D EPI protocols in terms of sensitivity and specificity to stimulus-related activation.

## Methods

### Image Acquisition

The cervical spinal cords (C4-C7 vertebral levels, approximately corresponding to spinal cord segments C5-C8) of eleven healthy volunteers (ages 22-39, 5 males, 6 females) were scanned using a 7 T whole-body MRI system (Magnetom 7 T AS, Siemens Healthineers, Erlangen, Germany) equipped with a G_max_ = 70 mT/m and maximum slew rate = 200 mT/m/ms gradient coil and a single-channel four-element transmit, 22-channel receive array brainstem and cervical spine radiofrequency (RF) coil^47^. All imaging experiments were performed under an IRB-approved protocol with subject consent and in full compliance with all applicable human subject protection regulations. Shim adjustment (up to 2^nd^ order) was performed during free breathing. B_1_^+^ adjustment was performed by setting transmit reference voltage to 250 V, a level which avoids clipping of RF pulses, and produces correctly-calibrated RF pulses in most subjects at approximately the level of the C5/6 intervertebral disc. Anatomical T_1_-weighted sagittal MP2RAGE^48^ images at 0.7 mm isotropic resolution were acquired for every subject to aid in registration to template space.

One single-shot 2D EPI protocol at 0.75 mm in-plane resolution and two multi-shot 2D EPI protocols at 0.75 mm and 0.6 mm in-plane resolution were used (Figure 1). These three protocols will be abbreviated SS75, MS75, and MS60, respectively. Each protocol was optimized independently, rather than attempting to match parameters between protocols. Research pulse sequences were used for both the single-shot and multi-shot acquisitions. (At the time of this study, a multi-shot 3D EPI pulse sequence analogous to what was used in references^12–14^ was not available on our scanner platform.) In all protocols, the FOV was intentionally centered towards the back of the neck to include posterior neck muscles, which, due to their greater distance from the trachea and esophagus, serve as spatial landmarks that are relatively robust to swallowing to aid subsequent motion correction.

In the single-shot EPI, a single saturation slab was placed anterior to the spinal cord for outer volume suppression, thereby reducing the required echo-train length. For in-plane acceleration using generalized autocalibrating partially parallel acquisition (GRAPPA)^49^, 52 lines of autocalibration signal reference data were acquired using GRE^50^, which has been shown to be optimal for single-shot EPI in the spinal cord at 7 T^51^. Partial Fourier (factor 6/8) acquisition was judged to be necessary in single-shot EPI to further reduce TE to avoid excessive signal loss due to through-slice dephasing. Image reconstruction of the single-shot EPI was performed online with the standard reconstruction pipeline of the manufacturer, including partial Fourier reconstruction by zero-filling and sine-window filtering, and Nyquist ghost correction based on a three-line navigator.

The multi-shot protocols were acquired without a spatial saturation slab due to SAR limitations given the shorter per-shot TR. Extending the volume acquisition time to accommodate spatial saturation was judged to be an unfavorable tradeoff. Therefore, a larger FOV had to be acquired in the multi-shot protocols, than in the single-shot protocol, to encompass signal from the entire neck. Note that phase-encode FOVs cannot be defined to be larger than readout-encode FOV, and the readout-encode FOV can be enlarged with no SNR or timing penalties. For this reason, the FOV in the multi-shot protocol was set larger in the readout (right-left) dimension than in the single-shot protocol to achieve the intended increase in the phase-encode FOV. Phase-encoding in the left-right dimension is undesirable because the patterns of spatial distortion in the spinal cord would be asymmetric across the left-right dimension of the spinal cord. Unlike the single-shot sequence, the custom multi-shot sequence did not allow for GRAPPA acceleration via the console, therefore the FOV in the phase-encode dimension was set to half of the readout FOV in the multi-shot protocols to achieve the intended phase-encode undersampling (Fig. 1). Image reconstruction of the multi-shot acquisitions was performed offline, with an iterative conjugate gradient SENSE algorithm^52^. The coil sensitivities were estimated based on a low-resolution gradient-echo acquisition. For SNR-optimal coil combination, the noise correlation between coils was calculated based on data acquired in vivo without RF excitation. A three-line navigator was acquired for each shot, which was used for per-shot frequency offset demodulation to reduce ghosting caused by temporal B_0_ field fluctuations^46^. The frequency offset per shot was estimated based on a magnitude-weighted coil combination of the phase at the center of k-space in the second navigator echo. For Nyquist ghost correction, three-line phase reference data (which characterize the gradient waveforms) were also acquired in a phantom immediately following each session, using the same multi-shot protocols as performed in vivo. In pilot studies, this approach was observed to be more effective in reducing Nyquist ghosting than using the three-line navigators from the in vivo data.

**Figure 1:**
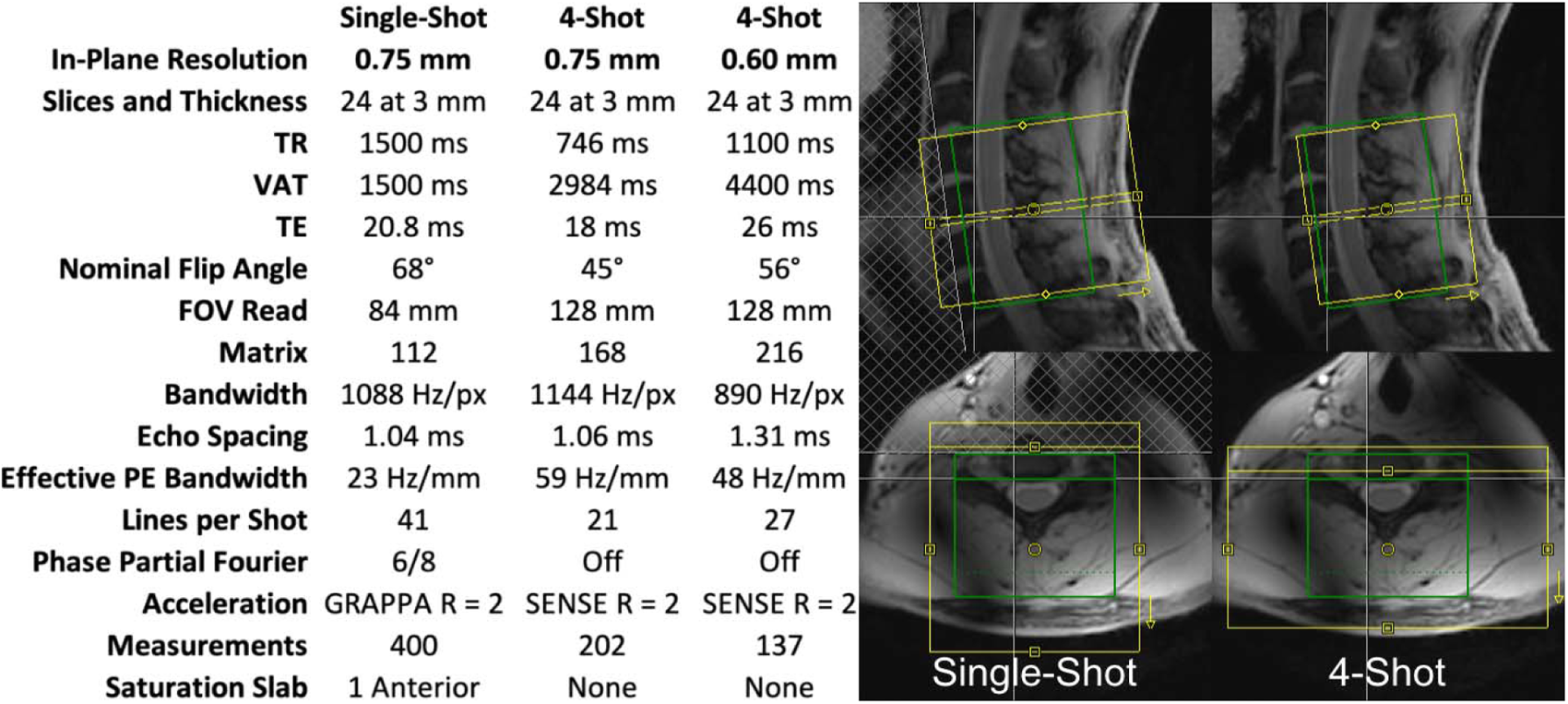
Pulse sequence parameters and slice positioning for three fMRI protocols. Yellow boxes indicate image acquisition fields of view, and green boxes indicate frequency and shim adjustment volumes.

To compare expected advantages and disadvantages between the protocols, Table 1 lists performance considerations relevant to BOLD fMRI, and for each of these gives the governing protocol parameters, and the expected relative ranking of the three candidate protocols. The relative importance of the different performance considerations, and the holistic ranking of protocol performance in cervical spinal cord BOLD fMRI at 7 T, is not obvious; the goal of this study is to determine this experimentally.

**Table 1:** Performance considerations relevant to BOLD fMRI, the protocol parameters governing performance regarding each image quality or sensitivity consideration, and the expected performance of the three candidate protocols relative to each other.

| Performance Consideration | Governing Parameter(s) | Expected Ranking |
| --- | --- | --- |
| BOLD Sensitivity | TE | **MS60** > SS75 > MS75 |
| Signal Dropout | TE | **MS75** > SS75 > MS60 |
| Ghosting | Number of Shots, TE | **SS75** > MS75 > MS60 |
| Partial Fourier Blurring | Partial Fourier Factor | **MS60** ≈ **MS75** > SS75 |
| T2* Decay Blurring | Effective Phase Encode Bandwidth (dk/dt) | **MS75** > MS60 > SS75 |
| Susceptibility-Induced Distortion | Effective Phase Encode Bandwidth (dk/dt) | **MS75** > MS60 > SS75 |

### Functional MRI Task Design

For each of the three imaging protocols, a 10-minute fMRI acquisition was performed with noxious thermal stimulation applied to the lateral surface of the base of the right thumb with a 30 mm x 30 mm flat thermode at a pre-calibrated intensity of 3/10 (range 42-51°C, average 46.5°C) using an fMRI-compatible thermal stimulator (TSA-II, Medoc, Israel). Pain scale anchors were 0 = no pain and 10 = worst pain imaginable. Stimulus blocks were 10 seconds in duration, calculated by including ramp-up and plateau, but excluding ramp-down to baseline. Ramp speed was set to 10°C/sec. Stimulus blocks were separated by pseudo-randomized inter-stimulus intervals ranging from 25 to 45 seconds, plus initial and final stimulus-free blocks. This stimulus was expected to produce sensory activation primarily in the ipsilateral dorsal horn at the neurological C6 (approximately vertebral C4-C5) level^53,54^. Subjects were coached to remain still and maintain steady respiration during thermal stimuli. Pulse-oximeter and respiratory traces were acquired (TSD200, PPG100C, TSD201, and RSP100C, Biopac, Goleta, CA, USA).

### Data Processing

Spinal cord toolbox v5.4.0^55^ was used for image pre-processing. Anatomical MP2RAGE images were segmented using *sct_deepseg_sc*, and vertebral levels were labeled using *sct_label_vertebrae* and/or *sct_label_utils*. Registration to the PAM50 template^56^ was performed using *sct_register_to_template*. Each 4-D fMRI dataset was motion-corrected using *sct_fmri_moco* with an inclusive cylindrical mask that contains the posterior neck muscle, the spinal canal and spinal cord were manually masked in FSLeyes^57^, and the fMRI dataset was registered to the PAM50 template using *sct_register_multimodal*, initialized using the transformation calculated between the anatomical image and PAM50. Each fMRI dataset was straightened, spatially smoothed using *sct_smooth_spinalcord* with an anisotropic 2 mm x 2 mm x 6 mm Gaussian kernel oriented along the spinal cord centerline, and unstraightened, to minimize blurring of CSF signal into the cord.

First-level GLM analyses was performed in individual-subject space using FSL FEAT^58^, incorporating the default autocorrelation correction using FSL FILM^58^ and a 35-term physiological noise model (PNM)^31,59^. The PNM was generated separately for each slice, incorporating slice timing correction, using FSL PNM, and included 8 cardiac phase, 8 respiratory phase, 16 cardiac-respiratory interaction, CSF signal derived from the spinal canal mask (canal mask minus cord mask), and two in-plane translational motion correction terms from *sct_fmri_moco*.

Group-level GLM analysis was performed in FSL FEAT using a mixed-effects model (FLAME1). After warping the results of first-level analyses into PAM50 template space, group-level activation maps were calculated within a search region constructed by eroding the PAM50 cord mask by one voxel. First, we performed thresholding at z > 2.7 followed by cluster-level correction at a cluster threshold of p = 0.05. This cluster-level correction method was tested because of its widespread use and familiarity in the field of brain fMRI, but has not yet been rigorously validated in the spinal cord, and so these results should be interpreted with caution. Next, because cluster-level correction did not yield any supra-threshold activation for any of the three protocols, we applied a low, uncorrected threshold of z > 1.64, which corresponds to a voxel p-value threshold of 0.05. This uncorrected threshold of z > 1.64 was subsequently used for tabulation of activated voxel counts and visualization in support of the goal of comparing the three acquisition protocols.

### Data Analysis

Temporal signal to noise ratio (tSNR) was calculated voxelwise as the ratio of temporal mean to standard deviation, then averaged within neurological level C6 in the PAM50 cord, gray matter, and white matter masks after motion correction, but before smoothing and application of the PNM. The extent of this neurological C6 mask was defined generously as the region where the probabilistic mask of C6 contained within SCT^60^ is greater than zero. Several additional ROI masks were defined for further analysis (Figure 2). The PAM50 cord mask, eroded by 1 voxel, was used as the search region in group-level analyses and to mask parameter maps for visualization. A mask for expected true-positive activation was constructed by summing the PAM50 masks for the right dorsal horn, right fasciculus gracilis, right fasciculus cuneatus, and right lateral corticospinal tract (PAM50 atlas mask entries 1, 3, 5, and 35), binarizing, eroding by one voxel from the cord-CSF border only, and calculating the intersection of the result with the aforementioned PAM50 mask for spinal (i.e., neurological) level C6 thresholded at zero. White matter structures were included to enclose potential BOLD signal in veins draining the expected gray matter site of activation^61^, but the spinal venous plexus on the exterior of the spinal cord, though expected to also exhibit BOLD activation^62^, was excluded due to the strong effects of CSF signal that cannot be fully accounted for by a PNM and the resulting strong potential for false-positive activation. A control mask, for comparison purposes, was calculated similarly using PAM50 masks for the left lateral corticospinal tract, left rubrospinal tract, left spinothalamic and spinoreticular tracts, and left ventral horn (PAM50 mask entries 4, 8, 12, and 30). These masks were also warped to each individual subject space. Although studies in rats have shown that some contralateral ventral horn activation does occur in response to noxious stimuli^63,64^, this was judged to be the most suitable control region in light of the stronger resting-state connectivity between ipsilateral dorsal and ventral horns, and between ipsilateral and contralateral dorsal horns, in humans^12,13,65^.

**Figure 2:**
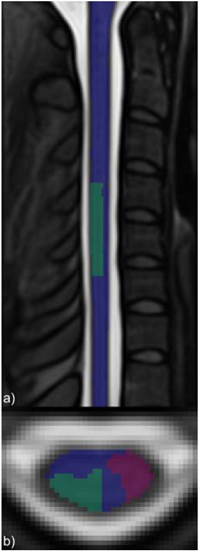
Masks used for analysis of task fMRI activation, displayed in PAM50 template space. The PAM50 cord mask, eroded by 1 voxel (blue), was used as the search region in group-level analyses, and to mask all parameter maps for visualization. Masks representing regions of interest in the C6 spinal level for expected true positive activation (green) and control (red) are also displayed.

For the group-level and each individual-subject result, the mean and maximum z-statistic across voxels with z > 1.64 within the true-positive and control ROI masks were tabulated. The ratio of the maximum z-statistic in the true-positive ROI mask to the maximum z-statistic in the control ROI mask in each subject was taken as a measure of the specificity of each protocol to true-positive activation. Mean BOLD percent signal change (PSC) was calculated in activated native space voxels in the true-positive region for each subject, and the BOLD percent signal change in the voxel in the true-positive region having the highest z-statistic in each subject was also tabulated.

## Results

Individual subject-level mean and single-timeframe images and tSNR maps of datasets representing the best-case and worst-case image quality in the study are shown in Figure 3. These best-case and worst-case datasets were chosen by first selecting the subsets of cases with the highest and lowest tSNR, and from those subsets, choosing the dataset with the lowest and highest artifact burden by visual inspection, respectively. In axial slices near intervertebral junctions, where residual B_0_ inhomogeneity is large, signal loss due to through-slice dephasing and geometric distortion are evident. Geometric distortion was less severe in multi-shot acquisitions, which had higher effective phase encoding bandwidth. The single-shot EPI had a blurrier appearance than the multi-shot acquisitions, while multi-shot 0.60 mm produced the sharpest appearing images. A spatial gradient in tSNR was also evident, with lower tSNR at more inferior levels. This is consistent with a combination of reduced B_1_ ^+^ efficiency, increased static B_0_ inhomogeneity, and larger time-varying B_0_ fluctuations due to the proximity to the lungs.

**Figure 3:**
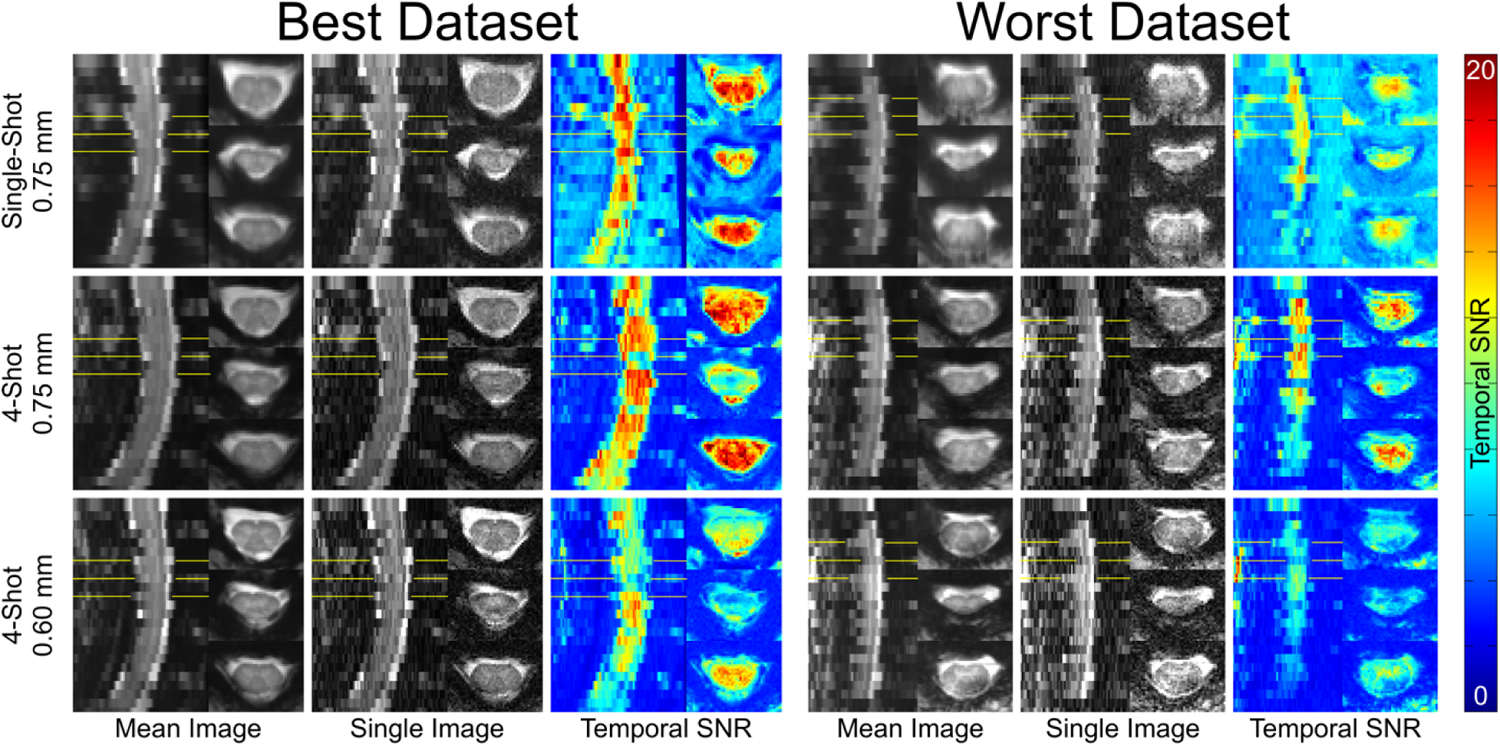
Individual subject-level images and temporal signal to noise ratio (tSNR) maps spanning C4-C7 vertebral levels in datasets representing best and worst image quality in the study. Temporal SNR was calculated after motion correction, but before spatial smoothing or application of the physiological noise model. Locations of the three axial slices are indicated with yellow lines in the sagittal frames. The first and third displayed axial slices are superior and inferior to an intervertebral disc, and the second axial slice is located at an intervertebral disc.

Summary statistics of tSNR are given in Table 2. Temporal SNR was generally highest in single-shot 0.75 mm and multi-shot 0.75 mm, with lower variability between subjects in multi-shot 0.75 mm, while tSNR was significantly lower in multi-shot 0.6 mm, yet with lowest between-subject variability. Temporal SNR was slightly greater in gray matter than in white matter in all three protocols, driven predominantly by the higher signal in gray matter.

**Table 2:** Summary statistics of temporal signal to noise ratio (tSNR) measurements in data from three protocols. Tabulated means and standard deviations are calculated as mean and standard deviation of the set of individual subject-level means within PAM50 masks. NB: standard deviations across subjects of mean tSNR in the cord, GM, and WM do indeed all round to 4.2; differences appear at the third decimal place.

| Temporal SNR in Neurological Level C6 |  |  |  |
| --- | --- | --- | --- |
|  | SS75 | MS75 | MS60 |
| tSNR in Cord | 14.6 ± 4.2 | 14.2 ± 3.3 | 10.0 ± 2.5 |
| tSNR in GM | 15.4 ± 4.2 | 15.1 ± 3.7 | 10.8 ± 2.6 |
| tSNR in WM | 14.2 ± 4.2 | 13.7 ± 3.0 | 9.7 ± 2.5 |

Using cluster-level correction for multiple comparisons within the eroded PAM50 cord mask, no significantly activated voxels were identified in the spinal cord on group level analyses for any of the three protocols. Maps of z-statistics, thresholded at an uncorrected threshold of z > 1.64, and BOLD percent signal changes in group-level GLM analyses are shown in Figure 4. Clusters of activation under this uncorrected threshold appear in all three protocols in the ipsilateral (right) dorsal quadrant at the expected C6 neurological level, which is centered approximately at the C4-C5 intervertebral disc (see Figure 2 for superior-inferior extent). All images are displayed in radiological orientation convention. The ipsilateral (right) activation appears in the same cross-sectional location in single-shot 0.75 mm and multi-shot 0.6 mm; however, this cluster is weaker and located more peripherally in multi-shot 0.75 mm. In single-shot 0.75 mm, a cluster also appears in the contralateral (left) dorsal horn.

**Figure 4:**
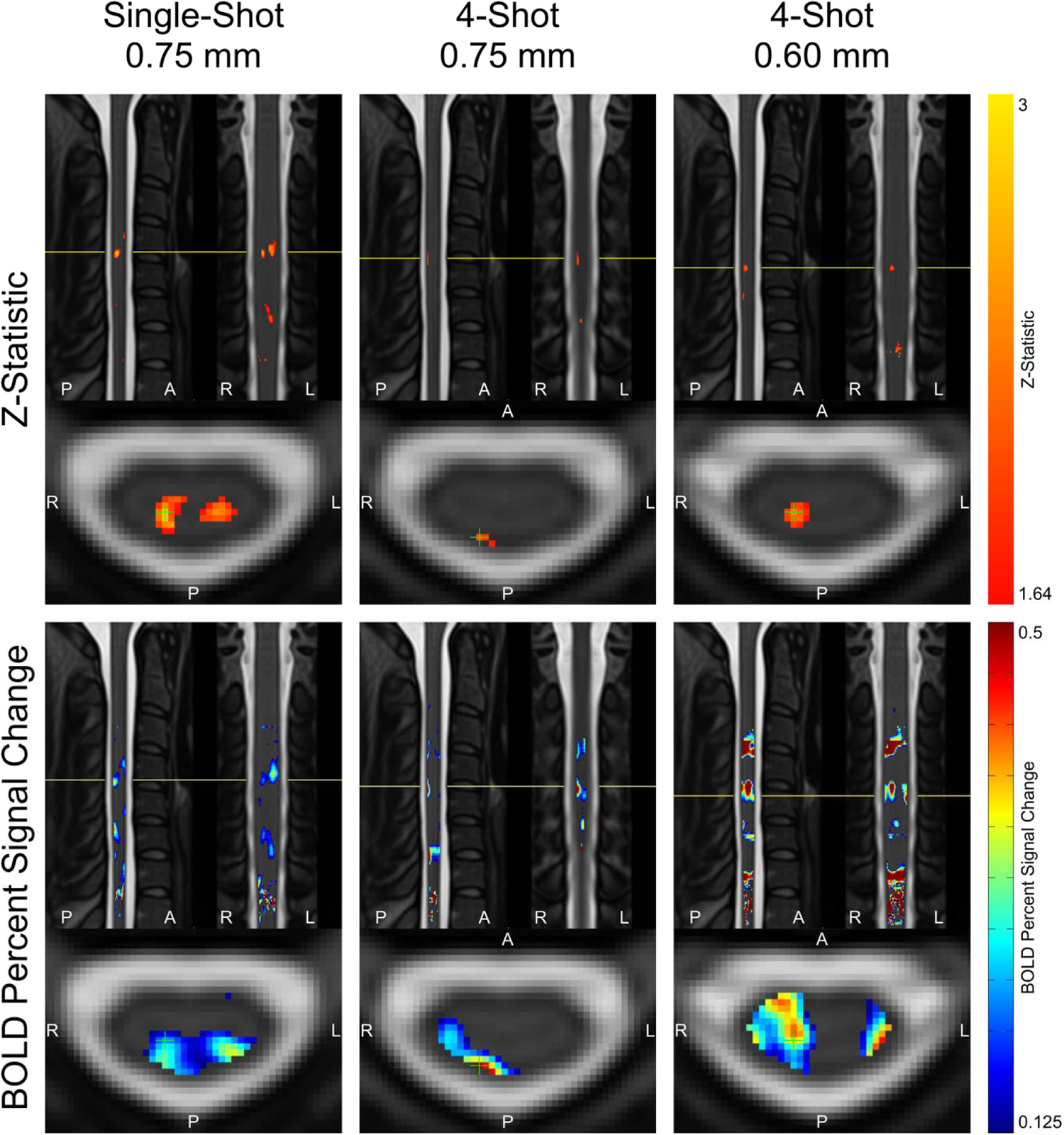
Maps of z-statistics and BOLD percent signal changes in group-level GLM analyses. Z-statistic maps are displayed using a low, uncorrected threshold of z > 1.64, and as such, should be interpreted with caution. Maps of BOLD percent signal change are thresholded at 0.125% for ease of visualization. Maps are masked to the spinal cord, encompassing all neurological levels, and the slice locations in all three planes are centered on the voxel inside the spinal cord mask with the highest z-statistic (indicated with a green crosshair).

Details of z-statistics and BOLD percent signal change in group-level GLM analyses are given in Table 3. Voxels within the true-positive and control ROI masks with z > 1.64 are considered “activated” for the purpose of comparisons between protocols, although as stated previously, these tabulated voxels do not survive multiple-comparison correction using cluster-level correction. Mean z-statistics in activated true-positive voxels are relatively consistent in the range of 1.83 to 2.04 across the three protocols, with single-shot 0.75 mm yielding the highest mean z-statistic contrary to initial expectation (Table 1). The maximum true-positive z-statistic is highest in single-shot 0.75 mm at 2.85, compared to 2.07 in multi-shot 0.75 mm and 2.38 in multi-shot 0.6 mm. All three protocols yielded a ratio of true-positive to control max z-statistic between 1.27 and 1.37.

The mean BOLD percent signal change in activated voxels was higher in multi-shot datasets than in single-shot. Multi-shot 0.75 mm yielded the highest mean BOLD percent signal change, at 0.43%, and the BOLD percent signal change in the voxel with the highest z-statistic was 0.50%.

**Table 3:** Summary statistics of group-level GLM results. An uncorrected threshold of z > 1.64 was used to tabulate activated voxel counts for the purpose of comparing acquisition protocols; therefore, these results should be interpreted with caution.

| Group-Level Activation Results |  |  |  |
| --- | --- | --- | --- |
| Threshold for Significance is $Z > 1.64$ | | | |
|  | SS75 | MS75 | MS60 |
| Mean Z-Stat in Activated Voxels in True Positive Region | 2.04 | 1.83 | 1.84 |
| Max Z-Stat in True Positive Region | 2.85 | 2.07 | 2.38 |
| Max Z-Stat in Control Region | 2.24 | 1.51 | 1.83 |
| Ratio of Max Z-Stat in True Positive vs. Control | 1.27 | 1.37 | 1.3 |
| % of Voxels Activated in True Positive Region | 2.76% | 0.71% | 2.34% |
| % of Voxels Activated in Control Region | 2.20% | 0.00% | 0.37% |
| Ratio of % of Voxels Activated in True Positive vs. Control | 1.25 | ∞ | 6.36 |
| Mean BOLD PSC in Activated Voxels | 0.21 | 0.43 | 0.33 |
| BOLD PSC in Voxel w/ Highest Z-Stat | 0.22 | 0.5 | 0.38 |
| Mean BOLD PSC in Voxels w/ Top 10 %ile Z-Stats | 0.19 | 0.26 | 0.33 |

To estimate the number of required subjects to achieve a supra-threshold cluster using stringent cluster-level correction methods, we performed additional group-level analyses with cluster-level correction as described in the methods section, on datasets containing replicates of the set of 11 individual subject-level datasets. Based on these analyses, we estimate that using thresholding at z > 2.7 followed by cluster-level correction at a cluster threshold of p = 0.05, the required number of subjects is estimated to be between 23 and 33 in single-shot 0.75 mm, between 177 and 187 in multi-shot 0.75 mm, and between 34 and 44 in multi-shot 0.60 mm.

Details of z-statistics and BOLD percent signal change across all individual-level GLM analyses are given in Table 4. As in group-level analyses, voxels within the true-positive and control ROI masks (warped to each individual subject space) with z > 1.64 are considered activated. Mean z-statistics in activated true-positive voxels are relatively consistent in the range of 1.96 to 2.18 across the three protocols. Single-shot 0.75 mm yields the highest mean z-statistic, but the standard deviation of each subject’s mean z-statistic in activated true-positive voxels is double that of the multi-shot protocols. Single-shot 0.75 mm also yields the highest mean maximum z-statistic at 2.68, compared to 2.27 for multi-shot 0.75 mm and 2.45 for multi-shot 0.6 mm. Single-shot 0.75 mm and multi-shot 0.6 mm have true-positive-to-control ratios substantially greater than unity, while the ratio in multi-shot 0.75 mm is closer to unity.

Similar to group-level analyses, the mean BOLD percent signal change in activated voxels was higher in multi-shot protocols. Multi-shot 0.6 mm yielded the highest mean BOLD percent signal change, at 1.78% ± 0.89%, followed by multi-shot 0.75 mm at 1.29% ± 0.76%, and single-shot 0.75 mm at 0.67% ± 0.31%. The same relationship among the protocols is evident in the BOLD percent signal change in the voxel with the highest z-statistic in each dataset.

**Table 4:**
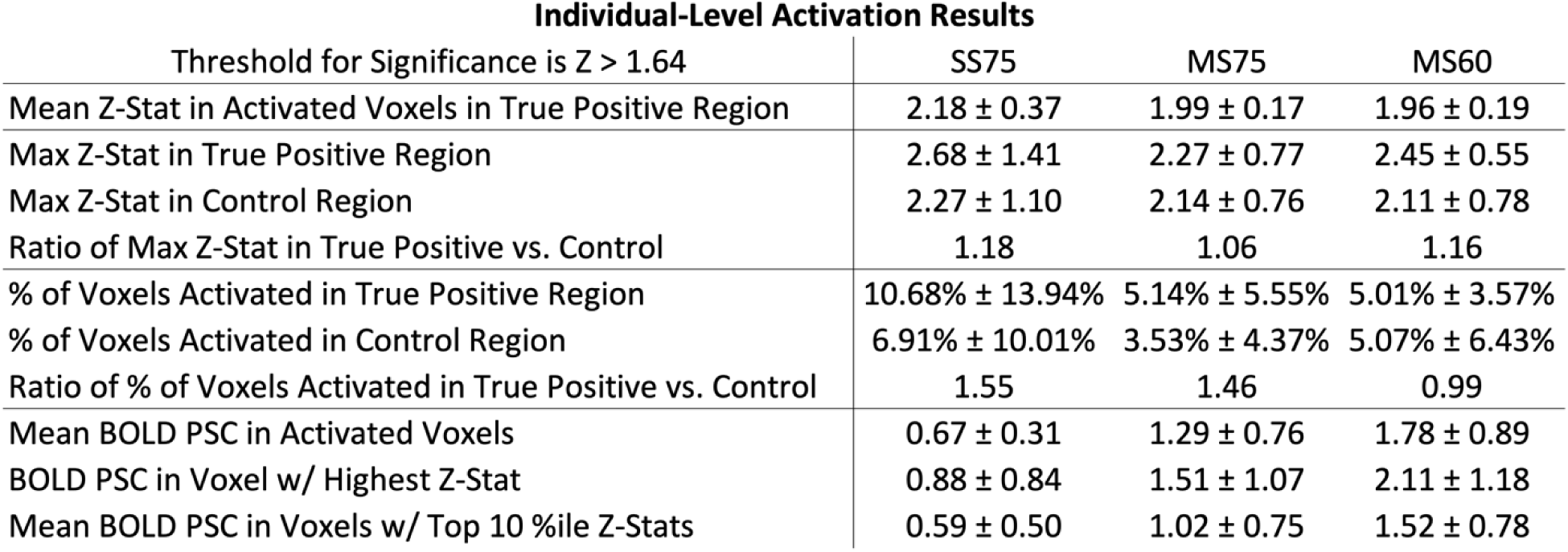
Summary statistics of individual subject GLM analyses. An uncorrected threshold of z > 1.64 was used to tabulate activated voxel counts for the purpose of comparing acquisition protocols; therefore, these results should be interpreted with caution. In individual subjects, mean z-statistics and mean BOLD percent signal change (PSC) are calculated within voxels having z > 1.64, as well as BOLD PSC in the voxel having the highest z-statistic and in the set of voxels having z-statistics in the top 10^th^ percentile. Maximum z-statistics in true positive and control regions are also calculated in individual subjects without a threshold on z-statistic. Means and standard deviations across these individual-subject mean and maximum values are then calculated and tabulated here.

## Discussion

Image quality and tSNR are strongly impaired in spinal cord fMRI by static focal B_0_ inhomogeneity near the intervertebral junctions and temporally variable B_0_ inhomogeneity arising from the lungs. These two sources of B_0_ inhomogeneity were clearly evident in the images and tSNR maps shown in Figure 3. Near the intervertebral junctions, tSNR was sharply reduced and geometric distortion and signal loss were markedly more severe than in slices further from a junction. The gradual reduction in tSNR in lower slices was partially due to static and dynamic B_0_ inhomogeneity from the lungs, but reduced B_1_^+^ efficiency in the lower cervical spine was also a significant contributor^47^; signal intensity in mean and single images was lower adjacent to vertebral level C7, compared to C4-C6, even in the best-quality dataset.

In general, the single-shot 0.75 mm protocol provided the greatest sensitivity (albeit highest between-subject variability in z-statistics) to the expected activation, but also appeared to have the lowest effective in-plane resolution and largest geometric distortion. This was in line with expectations (Table 1), as the lower effective phase-encode bandwidth makes it more susceptible to T_2_* blurring and geometric distortion due to static B_0_ inhomogeneity, on top of blurring due to partial Fourier acquisition. Multi-shot 0.6 mm produced the most well-localized activation clusters, and the sharpest images, while being qualitatively less affected by geometric distortion caused by static B_0_ inhomogeneity than single-shot 0.75 mm. However, sensitivity to activation was reduced due to its longer VAT, which reduced the number of time points in an experiment of fixed duration, and the lower tSNR, both of which reduce the statistical power. Multi-shot 0.75 mm generally performed poorly; it neither showed the sensitivity to activation of the single-shot 0.75 mm protocol nor the spatial specificity of the multi-shot 0.60 mm protocol, as if sharing the drawbacks of each of the other protocols, without enjoying the benefits of either protocol to a similar degree.

Decades of results in the brain have consistently shown that longer TEs (i.e., close to the T_2_* of gray matter) are associated with greater sensitivity to activation. In the spinal cord at 7 T, however, the detrimental effects of B_0_ inhomogeneity severely impair data quality at longer TEs. A balance must be struck between image quality, which favors shorter TE, and sensitivity to BOLD signal, which favors TE close to T_2_* of gray matter, which is 29.3 ± 4.5 ms at 7 T^66^. The optimal TE for a given spinal cord fMRI pulse sequence will therefore likely be shorter than the T_2_* of gray matter.

In group-level GLM analyses, single-shot 0.75 mm and multi-shot 0.6 mm appeared to yield similarly-sized activation clusters, and the maximum z-statistic in single-shot 0.75 mm was substantially greater than the maximum z-statistic in multi-shot 0.6 mm (see Figure 4). In individual-level results, however, multi-shot 0.6 mm generally yielded more sharply-defined activation clusters in the ipsilateral dorsal quadrant (see Figure S1), and the difference between the maximum z-statistic in single-shot 0.75 mm and the maximum z-statistic in multi-shot 0.6 mm was much smaller than in the group level analysis. A possible explanation that unifies these observations is that multi-shot 0.6 mm was better able to spatially resolve activation in individual subjects, resulting in small clusters; however, when multiple subjects’ data was combined, the inherent physiological variability in the location of activation among different subjects^60^, as well as potential limitations in group-level registration of distorted spinal cord EPI images, resulted in these sharply-defined individual-level activation clusters spatially coinciding less fully in multi-shot 0.6 mm than did the broader activation clusters in single-shot 0.75 mm. These broader activation clusters in single-shot EPI may have arisen due to its greater statistical power, having more temporal frames due to a shorter volume acquisition time and higher tSNR due to the lower effective resolution.

As stated previously, the noxious thermal stimulus was expected to produce sensory activation in the ipsilateral dorsal horn at the neurological C6 (approximately vertebral C4-C5) level based on the fact that the majority of first-order neurons in the lateral spinothalamic tract synapse in the ipsilateral dorsal horn^53,54^. Indeed, virtually all previous spinal cord fMRI studies of noxious thermal stimulation at lower field strengths or in animals have observed the strongest activation in the ipsilateral dorsal horn. However, the lateral spinothalamic tract does send some projections to mainly interneurons in the contralateral hemicord as well^67–70^, so it is highly likely that relatively weaker activation observed in the contralateral dorsal horn is not spurious but is, in fact, physiologically plausible and expected, and this has been observed in prior spinal cord _fMRI work_^15,18,71–74^.

Based on TE, BOLD percent signal change would be expected to be highest in multi-shot 0.60 mm, followed by single-shot 0.75 mm and then multi-shot 0.75 mm. In individual-level results, multi-shot 0.60 mm did indeed yield the highest BOLD percent signal change. However, of the remaining two protocols, multi-shot 0.75 mm yielded higher BOLD percent signal change than single-shot 0.75 mm. One potential explanation for this is that the effect of spatial resolution outweighed the effect of the magnitude of the underlying BOLD activation in these individual-level measurements; the multi-shot 0.75 mm acquisition is expected to better resolve BOLD activation than the single-shot 0.75 mm acquisition. An alternative explanation specifically concerning the difference in the mean BOLD percent signal change in activated voxels between the multi-shot 0.75 mm and single-shot 0.75 mm acquisitions is that the single-shot 0.75 mm acquisition, which contained 400 temporal frames, was statistically better powered to detect low-BOLD-percent-signal-change activation than the multi-shot 0.75 mm acquisition, which contained only 202 temporal frames and may have been additionally hampered by greater physiologic noise contributions (e.g., ghosting). In other words, a voxel with a low BOLD percent signal change is more likely to be identified as significantly activated using a protocol with a larger number of temporal frames and less susceptibility to ghosting. This would result in a greater number of low-BOLD-percent-signal-change voxels being included in the set of significantly activated voxels in the single-shot 0.75 mm acquisition, decreasing the mean BOLD percent signal change calculated for that set of voxels. However, when the mean BOLD percent signal change is calculated for each protocol in the set of voxels having z-statistics in the top 10^th^ percentile within each individual subject, BOLD percent signal change remains lower in the single-shot 0.75 mm acquisition than in the multi-shot 0.75 mm acquisition. In the same percentile-based set of voxels in group-level results, single-shot 0.75 mm likewise has the lowest mean BOLD percent signal change. In light of these results, the first potential explanation based on spatial resolution may be more likely.

The superior-inferior location of peak activation was similar across the three protocols in group-level analyses, and appeared at the expected location in the center of neurological C6 (determined from the probabilistic spinal levels from the PAM50 atlas), based on the anatomical location of the thermal stimulus. The superior-inferior location of the peak activation was expected to vary across subjects due to natural physiological variation^60^, but was not expected to vary among the three protocols within each subject. Contrary to this expectation, we did observe substantial variation in the superior-inferior location of peak activation of up to 1 ½ vertebral levels across the three protocols within individual subjects (see Figure S1). The cause for this variation is unclear without conducting a reproducibility study by repeating all three protocols on the same participants. If the true underlying stimulus-related activation occurs across a large spatial extent in the superior-inferior dimension, then random noise as well as variations in physiological noise may shift the center of the detected activation between acquisitions in an unsystematic way, as observed. In that case, increasing the sensitivity to detect BOLD responses, e.g. by acquiring for longer, should yield extended activation clusters. The true underlying activation is however unknown, and the data in this work cannot convincingly support any conclusion. From resting-state studies, we would expect networks of correlated activity stretching over approximately one vertebral level^65^. Task-activated networks, and patterns of nociceptive activation in particular may, however, have a broader spatial extent to resting-state networks due to the fact that primary afferent projections may ascend or descend in Lissauer’s tract before synapsing^67,75,76^. A large spatial extent of the activated networks in the superior-inferior dimension may even be a prerequisite to detect activation clusters in group-level analyses with relatively few subjects. If the activation were narrowly localized within one subject, with a variable center along the vertebral levels between subjects, the statistical power to detect activation at the group level would be much diluted. Furthermore, it has been shown in the cerebral cortex that significant variation in activation maps exists across sessions, even in a single subject^77^, and when evaluating activation based on thresholded single-subject activation maps, even consistent activation can appear highly variable when the significance of this activation is near the defined threshold for visualization^78^. Further studies would be needed to investigate the true extent of the activation, and the reproducibility of individual-level activation clusters.

Several groups have developed methods to address distortion and through-slice dephasing in single-shot GRE EPI at 3 T through slicewise dynamic shimming methods using either slice-specific gradient offsets^39^ or z-gradient refocusing blips^79,80^. These methods, applied to single-shot EPI, may be promising alternatives to multi-shot 2D or 3D EPI at 7 T but, at the time of this study, were not sufficiently refined to implement routinely at 7 T. Also 3D multi-shot EPI sequences suitable for implementation in fMRI experiments were not readily available at the time of this study, despite encouraging results demonstrating their robustness to B_0_ inhomogeneity in resting-state fMRI studies at 7 T^12,13^. Such sequences remain a highly promising option worthy of future investigation. This study also used different reconstruction methods for single-shot and multi-shot EPI: on-scanner GRAPPA and offline SENSE, respectively. An ideal solution would be to reconstruct single-shot EPI data offline using the same SENSE method as the multi-shot EPI data; however, due to data size and transfer rate limitations, we were not able to save raw data for single-shot fMRI for all subjects.

This study is limited by its small sample size of 11 healthy subjects. Because of this, no significantly activated voxels were identified in the spinal cord on group level analyses using cluster-level correction for multiple comparisons. To enable visual and quantitative comparisons between protocols, maps and tables used a low, uncorrected threshold of z > 1.64. Future studies in a larger number of subjects would be needed to achieve significance under appropriately stringent thresholding methods.

It should also be noted that the BOLD percent signal change maps displayed in Figures 4, S1, and S2 display the amplitude of the BOLD signal but do not provide information on the variability of the BOLD signal, and so they should be interpreted with appropriate caution, and in combination with the accompanying activation z-score maps.

## Conclusions

This study provides, to our knowledge, the first stimulus task fMRI results achieved in the spinal cord at 7 T. For a given scan time, single-shot 0.75 mm was most sensitive to the expected activation, likely due to its shorter volume acquisition time and greater statistical power, while multi-shot 0.60 mm provided the best-localized activation clusters and the least geometric distortion. The best choice of protocol will likely depend on the relative importance of sensitivity to activation versus spatial localization of activation for a given experiment.

## Acknowledgements

This study was supported by National Institutes of Health (NINDS) award K01NS105160 (ACS) and Department of Defense MSRP grant W81XWH-17-1-0204 (JX). FE is supported by the Max Planck Society and the European Research Council (under the European Union’s Horizon 2020 research and innovation programme; grant agreement No 758974). The work was also supported by the Wellcome Trust (Strategic Award 102645/Z/13/Z, Senior Research Fellowship 202788/Z/16/Z). The Wellcome Centre for Integrative Neuroimaging is supported by core funding from the Wellcome Trust (203139/Z/16/Z). Data in this study was acquired using the University of Minnesota Center for Magnetic Resonance Research (CMRR) Multiband EPI pulse sequence software package.

The authors thank Dr. David O’Connor for helpful guidance on sample size estimation under cluster-level correction.

Portions of this study in smaller numbers of subjects have previously been presented as conference abstracts^61,81^. The data that support the findings of this study are available from the corresponding author upon reasonable request.

## Required Statements

### Data availability statement

The data that support the findings of this study are available from the corresponding author upon reasonable request.

### Ethics approval statement and patient consent statement

All imaging experiments were performed under an IRB-approved protocol with subject consent and in full compliance with all applicable human subject protection regulations.

### Permission to reproduce material from other sources

Preliminary results from this manuscript have previously been presented in conference abstracts. No material contained within this submission are encumbered by any restrictions related to their previous publication.

### Clinical trial registration

This work does not meet the definition of a clinical trial.

## Supplementary Materials

### Individual Subject-Level Activation Maps

Maps of z-statistics and BOLD percent signal changes in two representative individual-level GLM analyses are shown in Figure S1 (best-quality dataset) and Figure S2 (worst-quality dataset). Z-statistic maps are thresholded at an uncorrected threshold of z > 1.64.

In the best-quality dataset in Figure S1, clusters of activation appear in the ipsilateral dorsal horn in all three protocols. The superior-inferior location of the peak activated cluster in the three protocols varies within this single subject to a greater extent than the expected size of a spinal segment in a single subject, although all three locations are contained within the group-level probabilistic PAM50 atlas entry for the C6 neurological level. In single-shot 0.75 mm, the cluster extends into the ipsilateral ventral horn as well, and a smaller cluster appears in the contralateral deep dorsal horn. In multi-shot 0.75, a disconnected cluster of weak activation also appears in the ipsilateral ventral horn. The cluster in multi-shot 0.6 mm is intense and well-localized, coinciding with a cluster of high BOLD percent signal change approaching 2%. In the displayed slices, one likely false-positive voxel is visible at the contralateral cord-CSF boundary in the multi-shot 0.60 mm, and the contralateral cluster in single-shot 0.75 mm is neither clearly true-positive nor false-positive, but for the purpose of this analysis, a portion of it falls within the control mask ROI, and is treated as such in descriptive statistics.

**Figure S1:**
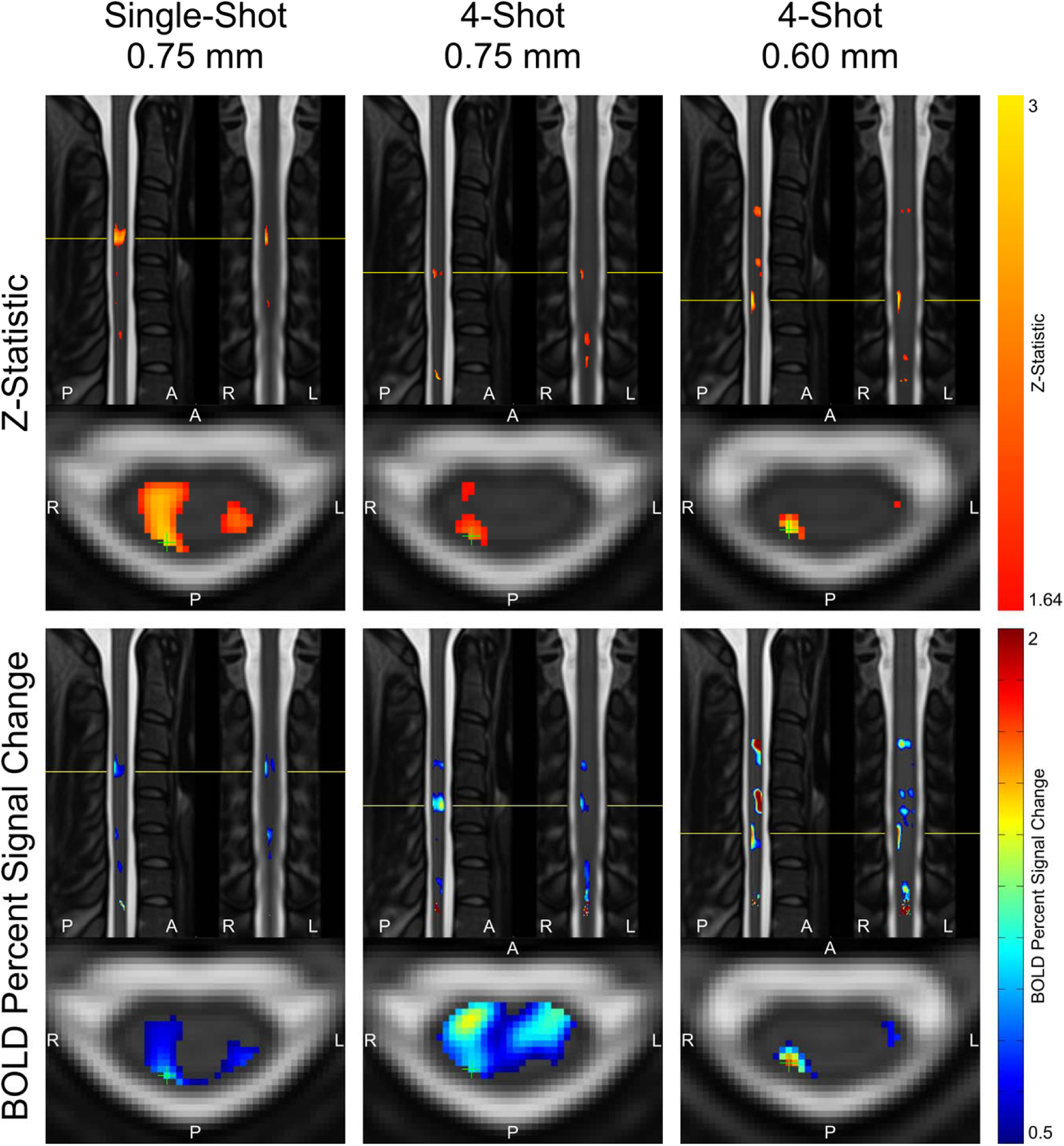
Individual subject-level maps of activation z-statistic and BOLD percent signal change, warped into template space, in the “best quality” dataset from Figure 2. Z-statistic maps are displayed using a low, uncorrected threshold of z > 1.64, and as such, should be interpreted with caution. Slice locations in all three planes are centered on the voxel with the highest z-statistic (indicated with a green crosshair).

In the worst-quality dataset in Figure S2, potentially plausible true-positive activation is visible in the ipsilateral dorsal quadrant in the multi-shot 0.6 mm protocol, though this cluster extends partially out of the superior boundary of the neurological C6 level. No activation exceeding the threshold of z = 1.64 is visible in either the single-shot or multi-shot 0.75 mm protocols. The location of the peak (although still sub-threshold) z-statistic within the true-positive mask ROI is indicated with a green crosshair in the axial plane, and yellow lines in the sagittal and coronal planes.

**Figure S2:**
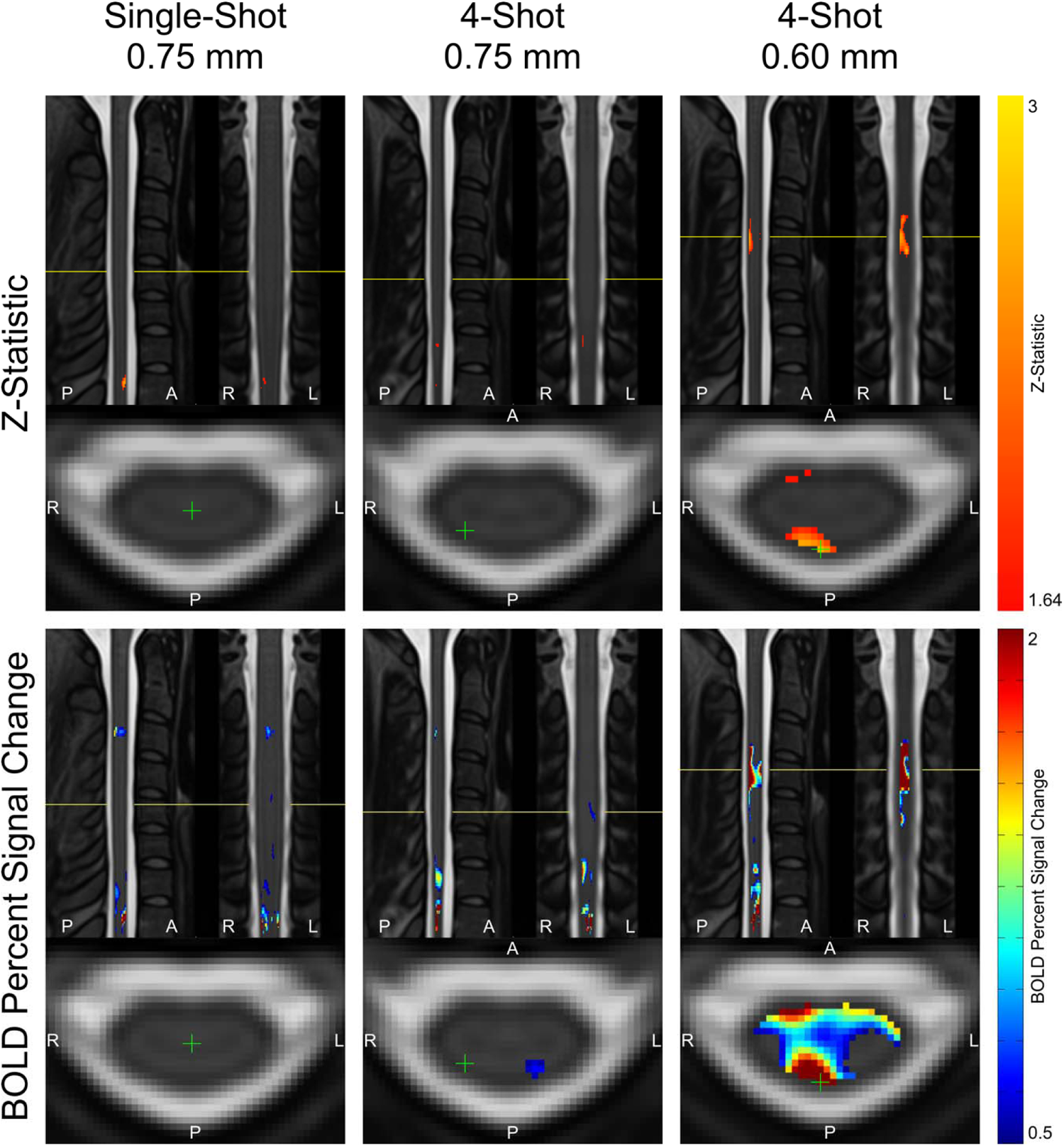
Individual subject-level activation maps, warped into template space, in the “worst quality” dataset from Figure 2. Z-statistic maps are displayed using a low, uncorrected threshold of z > 1.64, and as such, should be interpreted with caution. Images are centered at the voxel in the true positive region with the highest z-statistic (indicated with a green crosshair), but in single-shot 0.75 mm and 4-shot 0.75 mm, this z-statistic is below the threshold of 1.64.

